# Predicting drug response from single-cell expression profiles of tumours

**DOI:** 10.1101/2023.06.01.543212

**Authors:** Simona Pellecchia, Gaetano Viscido, Melania Franchini, Gennaro Gambardella

## Abstract

Drug response prediction at the single cell level is an emerging field of research that aims to improve the efficacy and precision of cancer treatments. Here, we introduce DREEP (Drug Response Estimation from single-cell Expression Profiles), a computational method that leverages publicly available pharmacogenomic screens and functional enrichment analysis to predict single cell drug sensitivity from transcriptomic data. We validated DREEP extensively *in vitro* using several independent single-cell datasets with over 200 cancer cell lines and showed its accuracy and robustness. Additionally, we also applied DREEP to molecularly barcoded breast cancer cells and identified drugs that can selectively target specific cell populations. DREEP provides an in-silico framework to prioritize drugs from single-cell transcriptional profiles of tumours and thus helps in designing personalized treatment strategies and accelerate drug repurposing studies. DREEP is available at https://github.com/gambalab/DREEP.

## Introduction

Intra-tumour heterogeneity (ITH) is a recurrent issue in designing effective treatment strategies in clinical practice. It refers to heterogeneous cancer cell populations present in the same tumour that exhibit high cellular plasticity and phenotypic heterogeneity. Accumulating evidence from independent studies shows that ITH gives rise to rare sub-populations of malignant cells characterized by a different epigenetic and transcriptional state that renders them refractory to a variety of anticancer drugs, protecting the whole population from complete eradication (1–4).

Single-cell RNA sequencing (scRNA-seq) has now become a powerful tool to explore ITH at the transcriptional level (5) and thus offers an unprecedented chance to tackle it therapeutically (4, 6, 7). In addition, large-scale pharmacogenomic screens have been performed across hundreds of molecularly characterized cancer cell lines (8–14). The resulting data have provided a valuable resource to link drug sensitivity with genomic and transcriptomic features. Indeed, in the last few years, several machine-learning models have leveraged these datasets to identify molecular markers linked to *in vitro* drug sensitivity in order to predict drug response from -omics profiles of tumour biopsies (8, 15, 16). However, these approaches do not explicitly account for ITH with the result that genomic and non-genomic biomarkers of drug response have been found for only a restricted number of small molecules. Nonetheless, expression-based biomarkers measured from bulk cell populations have emerged as the most powerful predictors of drug response *in vitro* for several cytotoxic and targeted drugs, with far more predictive power than other -omics profiles (8, 10). To overcome these limitations, a few recent studies like scDRUG (17, 18), scDEAL (19), beyondCell (20), chemCPA (21), and NCI-DOE (22) have attempted to leverage scRNA-seq and publicly available drug sensitivity profiles to predict drug response at the single cell level. However, these studies often exhibit certain drawbacks. They may lack comprehensive *in vitro* validation, fail to provide user-friendly software packages, rely on complex models requiring parameter fine-tuning before practical use, or are confined to predicting the efficacy of a limited number of drugs.

To address these challenges, we introduce DREEP (Drug Response Estimation from single-cell Expression Profiles), a novel bioinformatics tool designed to augment the precision and efficacy of therapeutic strategies by addressing the innate heterogeneity of cancer cell populations. DREEP’s primary function is to assess the complexity of tumour therapeutics and recommend specific drugs tailored to target distinct cell subpopulations. DREEP can predict a cell’s vulnerability to more than 2,000 drugs by simplifying the whole process with enrichment analysis, eliminating the need for complex parameter adjustments and thus making this type of analysis accessible to a broad audience.

We validate DREEP performance extensively using *in vitro* drug response data of over 200 cancer cell lines covering 22 distinct cancer types from publicly available single-cell and drug viability datasets. We show that our method can effectively identify recurring drug vulnerabilities across cells of diverse tumour types and establishes a clear relationship between drug response and the functional status of the cells. We show that DREEP can detect changes in drug sensitivity over time in the breast cancer cell line MCF7 and predict the emergence of drug resistance. Additionally, using molecular barcoded MDA-MB-468 cells (4), we show that our method can recognize innate drug-tolerant cell subpopulations and predict drug co-treatments that will inhibit all of them. Finally, we demonstrate that DREEP can accurately predict drug sensitivity within heterogeneous cancer cell populations of patient-derived cultures (PDCs) containing a mix of primary and metastatic cells derived from individuals with head and neck squamous cell carcinomas (23). In conclusion, our results demonstrate that it is possible to detect differences in drug response among individual cells within the same tumour. DREEP is implemented as an open-source R package, making it easily accessible and user-friendly for the broader scientific community. It is publicly available on GitHub at the following address https://github.com/gambalab/DREEP.

## RESULTS

### 1. Drug Response Estimation from single-cell Expression Profiles

DREEP (Drug Response Estimation from single-cell Expression Profiles) is a bioinformatics tool that leverages results from large-scale cell-line viability screens and enrichment analysis to predict drug vulnerability from the transcriptional state of a cell. It requires a *pre-defined* collection of Genomic Profiles of Drug Sensitivity (GPDS); these are lists of genes ranked according to their contribution in correctly predicting the effect of a small molecule. To build GPDS, we integrated several publicly available RNA-seq datasets and cell-line viability screens, including the following datasets: GDSC (8), CTRP2 (13, 24) and PRISM (25) (Figure 1A).

**Figure 1.**
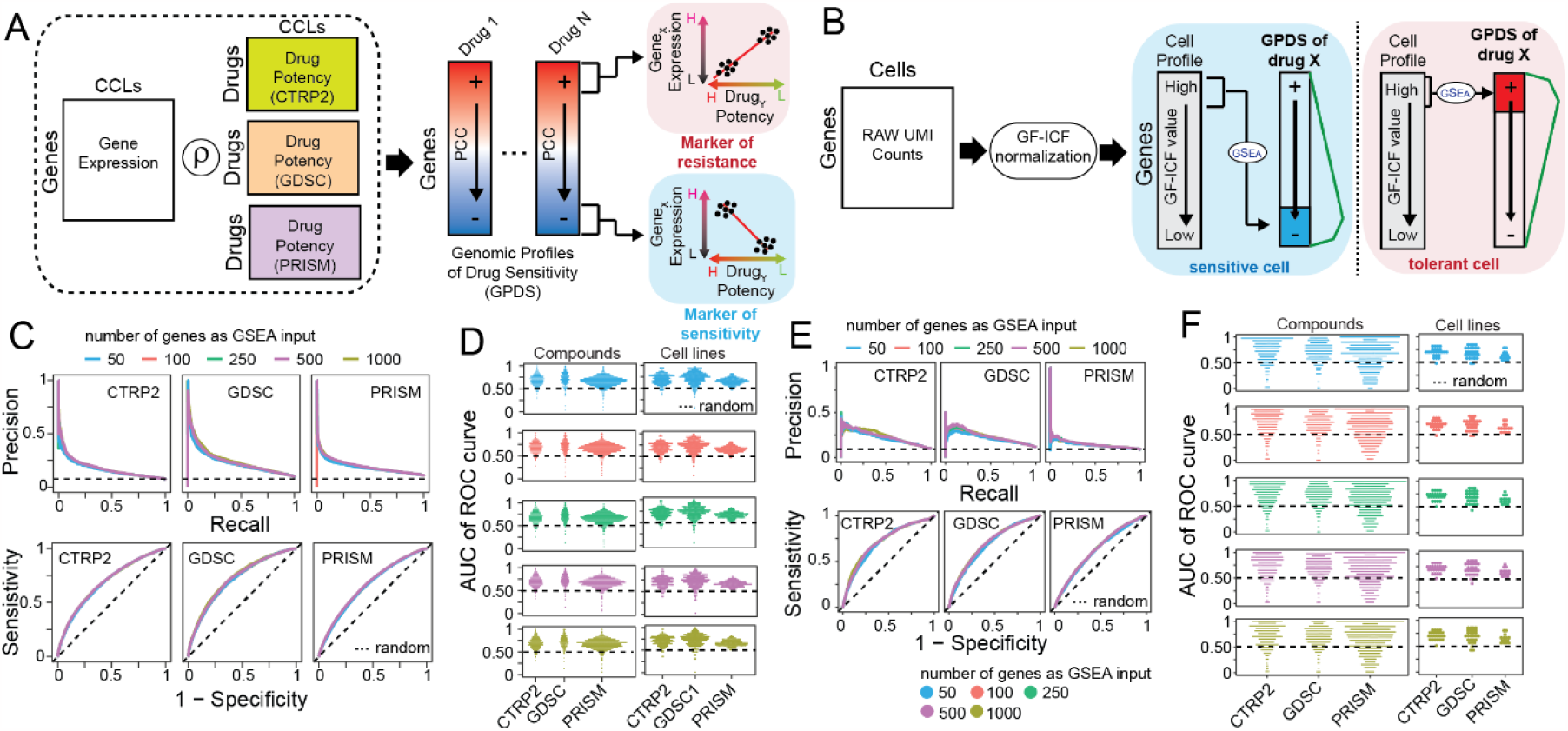
Drug Response Estimation from single-cell Expression Profiles (DREEP). (**A**) Construction of Genomic Profiles of Drug Sensitivity (GPDS). The potency profile of each drug in terms of AUC (Area Under the Curve) is correlated with the expression of all messenger RNA across the cell lines for which the drug potency was measured. (**B**) Bioinformatics pipeline for the prediction of single-cell drug response. Raw UMI counts are first normalized with the gf-icf pipeline and then the most relevant genes in a single cell are used as input for a GSEA against the GPDS ranked-list to predict its drug sensitivity. (**C**) Performance of DREEP drug sensitivity on 198 cell lines for each set of GPDS signatures used and different numbers of relevant genes to predict the effect of a drug. DREEP’s performance is estimated using either Precision-recall curve (upper) or ROC curve (bottom). Max AUC of ROC curves are CTRP2 = 0.691, GDSC = 0.728 and PRISM = 0.661. (**D**) Left column: Each point is a drug whose y-coordinate is AUC of the corresponding ROC curve across the 198 cell lines and computed by sorting cell lines from the most to the least sensitive to the drug. Each panel reports the results using different numbers of relevant genes to predict the effect of the drug. Right column: Each point is a cell line whose y-coordinate is the AUC of the corresponding ROC curve obtained by sorting drug predictions from the most to the least sensitive. Each panel reports the results using different numbers of relevant genes to predict the effect of a drug. (**E**) Same as (C) but for the 32 breast cancer cell lines published in Gambardella et al. (5). Max AUC of ROC curves are CTRP2 = 0.721, GDSC = 0.699 and PRISM = 0.637. (**F**) Same as (D) but for 32 breast cancer cell lines published in Gambardella et al. (5).

GPDS were computed by correlating the *in vitro* response to a small molecule from the above studies with the expression of all messenger RNA (mRNA) across all the cancer cell lines for which the drug potency was measured (Figure 1A and methods). In these studies, RNA-seq represents the basal gene expression of a cell line, while the potency of a small molecule in a cell line is evaluated as the Area Under the Curve (AUC) of the corresponding dose-response curve. The AUC value reflects cell growth inhibition after 72 hours of treatment, where lower values indicate sensitivity while higher values imply resistance to the tested drug. Thus, a gene whose expression positively correlates with the AUC of a drug across many cell lines can be considered predictive of drug resistance (i.e., the more expressed the gene, the higher the concentration needed to inhibit cell growth) (24). *Vice versa*, a negatively correlated gene can be considered predictive of drug sensitivity (*i*.*e*. the more expressed the gene, the lower the concentration needed to inhibit cell growth) (24).

Using this approach, we generated a collection of 2,434 GPDS signatures (i.e. 478 from GDSC, 511 from CTRP, and 1,445 from PRISM) and used them to predict the single-cell-level drug response with the strategy depicted in Figure 1B. Briefly, single cell RNA sequencing data from a sample was first normalized with the gf-icf (Gene Frequency – Inverse Cell Frequency) pipeline to rank genes in each cell according to their relevance (26–28). Then, for each cell we used the most relevant genes as input for Gene Set Enrichment Analysis (GSEA) (29) against each GPDS ranked-list. Since resistance biomarkers are at the top and sensitivity biomarkers are at the bottom of GPDS ranked lists, a positive value of the enrichment score (ES) returned by DREEP implies that the cell expresses genes associated with drug resistance. *Vice versa*, a negative ES indicates the cell expresses genes correlated with drug sensitivity. At the end of this process, it is thus possible to estimate the overall effect of each drug on a cell population by estimating the median DREEP enrichment score on that population and compute the percentage of cells sensitive to the drug by simply counting the number of cells in the sample that are associated with a significant ES less than zero (methods).

We found 90 small molecules shared among GDSC, CTRP2, and PRISM datasets. To assess the consistency of GPDS ranked lists, we calculated Spearman Correlation Coefficients (SCC) between GPDS representations of the same drug but constructed from distinct drug viability datasets. Supplementary Figure 01 shows a substantial similarity in GPDS rankings of the same drug even if derived from two distinct datasets, surpassing random comparisons by several folds (methods).

### 2. Estimation of the predictive performance of DREEP

To validate the GPDS ranked list and demonstrate the reliability of our method as well as measure its predictive performance on *in vitro* data, we applied it to the Kinker et al. dataset (30) comprising 53,513 single cell transcriptional profiles from 198 tumorigenic cell lines and spanning 22 distinct cancer types. To evaluate DREEP performance, we first converted predictions from the single-cell level to the cell-line level by computing the median enrichment score for each drug (i.e. GPDS) across the cells of each individual cell line. Then, to minimize potential confounders, we evaluated the performance of the DREEP method for each sensitivity dataset independently (Methods). We also searched for the optimal number of relevant genes to use for enrichment analysis and predict the sensitivity of a cell (*i*.*e*., 50, 100, 250, 500, and 1,000 genes). Figure 1C shows the method’s overall performance in terms of Precision-Recall and ROC (Receiver Operating Characteristic) curves across 198 cancer cell lines for each drug sensitivity dataset independently (i.e., CTRP2, GDSC and PRISM). In this analysis, predicted drug/cell-line pairs are ranked according to the median enrichment score estimated by DREEP and the performance is evaluated against the corresponding gold standard (methods). Figure 1D reports the ROC curve’s AUC (Area Under the Curve) for all drugs and all cell lines individually. Next, to investigate whether DREEP exhibited a prediction bias toward a class of drugs characterized by specific Mechanism of Action (MoA), we categorized the ROC curve’s AUC of each drug in Figure 1D (left panels) into two distinct groups: (i) Accurately Predicted (defined as ROC-AUC ≥ 0.75) and (ii) Erroneously Predicted drugs (defined as ROC-AUC ≤ 0.5, indicating random performance). This analysis yielded a total of 639 drugs whose efficacy was accurately predicted (209 from GDSC, 147 from CTRP2, and 282 from PRISM; Supplementary Table 01) and 135 drugs whose efficacy was erroneously predicted by DREEP (21 from GDSC, 23 from CTRP2, and 91 from PRISM; Supplementary Table 01). Notably, accurately predicted drugs exhibited a significant enrichment (31) (FDR < 10%) across a broad spectrum of MoAs (methods), spanning various biological mechanisms (Supplementary Table 02). In contrast, the 135 drugs predicted with low accuracy showed enrichment for a limited number of MoAs but overlapped with the MoAs of accurately predicted drugs (Supplementary Table 02), except for prostanoid and glucocorticoid agonist molecules. This analysis indicates that DREEP did not have a prediction bias toward a specific class of drugs, but it proved incapable of predicting the effects of these 135 small molecules, leading us to exclude them from the final version of the tool. Lastly, to assess DREEP’s ability to generalize across different cancer types, we grouped the ROC curve’s AUC values for individual cell lines (as reported in Figure 1D, right panels) according to their respective cancer types. Supplementary Figure 02 illustrates that DREEP does not reveal any significant performance drop associated with specific cancer types except for Neuroblastoma where average AUC is lower than in other cancer types. Finally, we conducted an additional assessment of DREEP’s predictive performance where we constructed GPDS ranked lists using the drug’s IC50 values instead of the AUC metric. As depicted in Supplementary Figure 03, DREEP’s performance noticeably decreased across all three drug viability datasets when employing this alternate approach.

To test DREEP’s performance on an additional independent dataset, we applied it to a single-cell breast cancer dataset (5). This dataset comprises 35,276 individual cells from 32 breast cancer cell lines covering all the main breast cancer tumour subtypes (Luminal/Her2-positive/Basal Like). Figure 1E,F show DREEP’s performance in this additional dataset.

Next, we tested the prediction performance of the DREEP method on both pan-cancer and breast cancer datasets using absolute gene expression levels instead of gf-icf normalized data. We observed a drastic reduction of the predictive performance of the method (Supplementary Figure 04) using the absolute gene expression level to estimate its relevance in a cell, showing the importance of data pre-processing prior to the application of a drug prediction algorithm. As a good compromise between the required computational time and prediction accuracy, we decided to use N = 500 genes as a default value for the DREEP sensitivity predictions in all subsequent analyses in the manuscript.

Finally, we used both the single-cell pan-cancer and breast cancer datasets to conduct a comparative analysis of DREEP’s performance with four other widely used single-cell drug prediction methods, including scDRUG (17, 18), scDEAL (19), beyondCell (20) (methods). We evaluated the performance of these methods based on the shared drugs, as depicted in Supplementary Figure 05A. As demonstrated in Supplementary Figures 05B-F, DREEP consistently outperformed the other methods across both datasets.

Taken together, these results collectively demonstrate the validity of our GPDS ranked lists for predicting the sensitivity of cells to drugs. Furthermore, our comprehensive evaluation showcases that DREEP consistently outperforms random predictions across various scenarios and outperforms other state-of-the-art methods.

### 3. The DREEP method reconstructs the therapeutic landscape of the pan-cancer dataset

Next, we used the DREEP method to reconstruct the therapeutic landscape of the 22 cancer types in the cell line pan-cancer dataset of Kinker et al. (30). As shown in Figure 2A,B (and Supplementary Table 03), we first used DREEP to convert each cell line into a list of small molecules predicted to be effective for at least 90% of profiled cells. We then grouped the cells and the drugs into four clusters each, so that cell lines that respond similarly across the tested drugs are grouped together, and analogously drugs that have an effect on the same cell lines are grouped together (i.e., cell line clusters C1, C2 C3, C4 and drug clusters D1, D2, D3 and D4).

**Figure 2.**
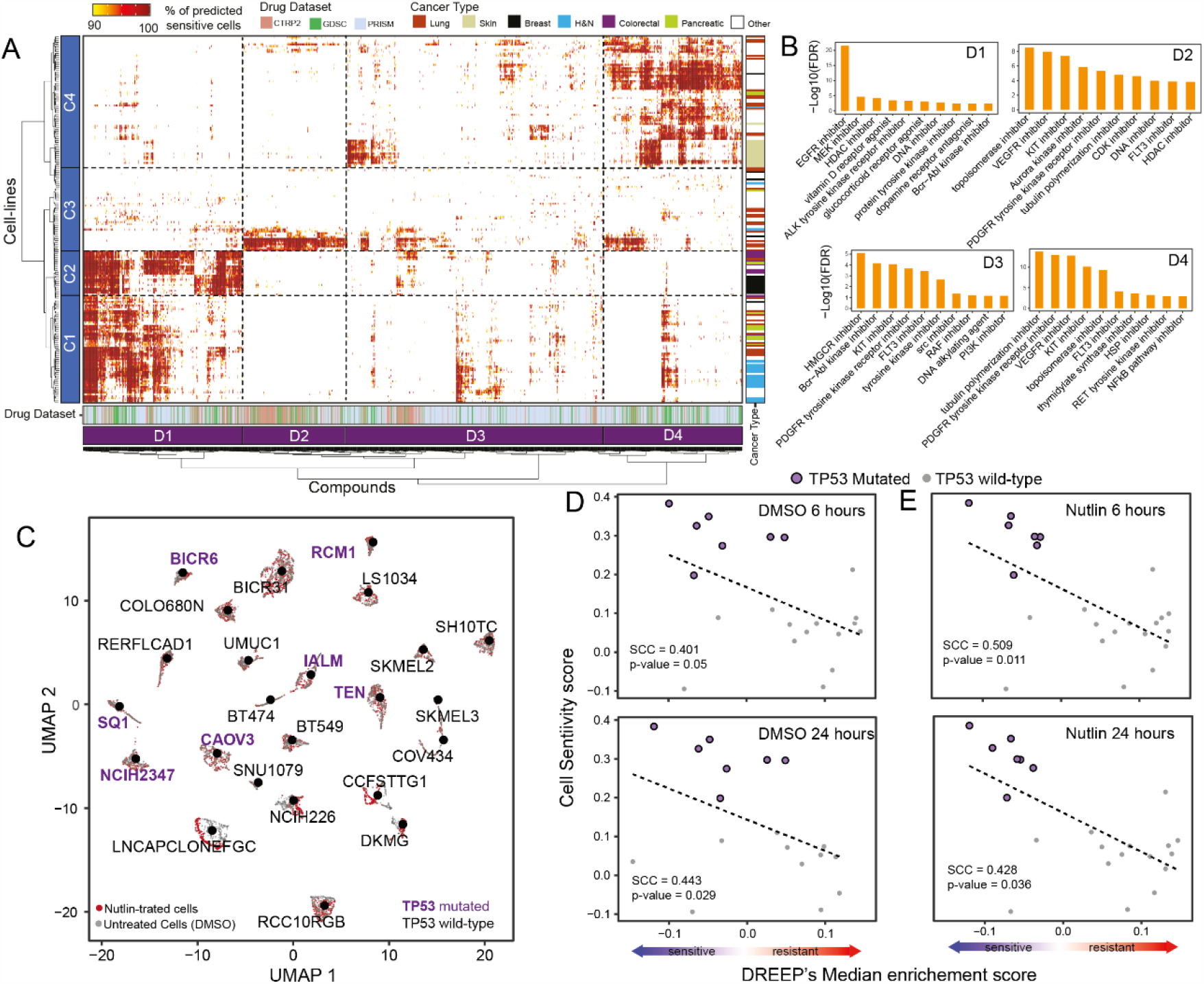
DREEP captures single cell variability in drug response. (**A**) Heatmap depicting both cell-line clusters (rows) and cancer-specific drug clusters (columns). Cell lines are colour-coded according to the cancer type while drugs are colour-coded according to the dataset to which they belong. Both rows and columns of the heatmap were grouped in four clusters named C1-C4 for cell line (rows) and D1-D4 for drugs (columns). (**B**) Bar-plot depicting enriched MoAs in each cancer-specific drug cluster. Only the top ten significant MoAs (FDR < 10%) for each cluster are shown. (**C**) UMAP plot of 7,805 cells of 24 cell lines either untreated or exposed to nutlin treatment. (**D**) The Spearman correlation coefficient (SCC) between observed and predicted sensitivity scores for the XX untreated (DMSO) cell lines in (C). In the upper plot the SCC is computed using cells sequenced after 6 hours, while in the bottom plot SCC is computed using cells collected after 24 hours. (**E**) Same as in (D) but for nutlin-treated cells.

These analyses revealed interesting associations between cancer vulnerabilities and drug response, highlighting numerous cell lines and cancer types predicted to be sensitive to small molecules with shared mechanisms of action (MoA) (31). Indeed, as shown in Figure 2A,B (and Supplementary Table 03 and Supplementary Table 04), cell lines in clusters C1 and C2 comprised 87% (13 out of 15) of head and neck cell lines, 100% of colorectal cell lines (10 out of 10), 83% of bladder cell lines (5 out of 6), 73% of breast cell lines (11 out of 15), 73% of pancreatic cell lines (8 out of 11), and 33% of lung cancer cell lines (13 out of 39). These cancers all have in common Epidermal Grow Factor (EGF) receptor overexpression/amplification, such as EGFR and ERBB2 (32). Indeed, this is one of the major alterations in colorectal (CRC, 60-80%) (33), lung (NSCLC, 40-80%) (34), head and neck (HNSCC, 80-90%) (35, 36), bladder (37, 38), pancreatic (PDAC, 30-89%) (39), and breast cancers (40). For these tumour types, therapies targeting EGFR, ERBB2, or their downstream effectors such as MEK or PI3K, have shown some degree of success (36, 41– 43). As depicted in Figure 2A,B, cell-line clusters C1 and C2 exhibit sensitivity to drugs in the D1 cluster that is enriched for drugs whose mode of action (methods and Supplementary Table 04) is related to the inhibition of EGFR and MEK signalling pathways.

Cluster C3 includes 69% of ovarian cancer cell lines (9 out of 13) and 70% of endometrial/uterine ones (7 out of 10). Cell lines in this cluster are predicted to be sensitive to drugs in the D2 cluster, which is enriched for topoisomerase, AURORA, HDAC, and CDK inhibitors. Interestingly, ovarian and uterine cancers are usually due to inefficient DNA repair (44). The standard of care for these cancer types is mainly based on neoadjuvant chemotherapy and several assays on cell lines identified drugs targeting apoptosis or cell cycle regulators as being effective (45, 46, 55–57, 47–54).

Cluster C4 includes all skin cancer cell lines (16 out of 16), all kidney (6 out of 6), all brain (12 out of 12), and 13 lung cancer cell lines. Cell lines in cluster C4 are predicted to be sensitive to multiple families of drugs, including RAF, KIT, PDGFR, and PI3K/mTOR inhibitors. This agrees with current clinical practice which uses mTOR and BRAF inhibitors as first-line therapy for kidney cancer and melanoma, respectively (58, 59), since the c-Met/mTOR axis is frequently altered in kidney cancer (60, 61) while BRAF mutations are the main genetic drivers of melanoma (59). Finally, RTKs are frequently altered in glioblastoma (62, 63).

To further demonstrate DREEP’s ability to identify recurring drug vulnerabilities across cancer cells of different tumour types with shared genetic backgrounds, we applied our method to the dataset from McFarland et al. (64). This dataset encompasses 3,630 untreated and 4,175 nutlin-treated cells, culminating in an extensive collection of 7,805 cells (Figure 2C). Cells were sampled at two time points, 6 and 24 hours post-treatment, and originated from 24 distinct cancer cell lines covering 14 different tumour types. Nutlin, a selective MDM2 inhibitor, is a negative regulator of the tumour-suppressor gene TP53, triggering apoptosis and cell cycle arrest exclusively in cancer cells retaining the wild-type TP53 form (65). As depicted in Figure 2C, seven of the 24 cell lines within this dataset harbour a missense mutation in the TP53 gene, while all other cell lines retain the wild-type (WT) TP53 form. Indeed, as demonstrated in Supplementary Figure 06, DREEP accurately predicted that most TP53 wild-type cells would exhibit sensitivity to nutlin treatment, while cells bearing TP53 mutations would display resistance. To validate these predictions, we first aggregated them at the cell line level by computing the median enrichment score of nutlin estimated by DREEP across individual cells within each respective cell line and then correlated these scores with the experimentally observed nutlin response from McFarland et al. (64) (methods). As depicted in Figure 2D,E, this correlation held true for both untreated and nutlin-perturbed cells and across the two time points. These findings further underscore the predictive accuracy of our method in effectively characterizing drug responses.

Altogether, these results show that DREEP can find recurrent drug sensitivity patterns shared by cell lines from multiple cancer types, as well as the relationship between drug response and functional cell status.

### 4. The DREEP method recapitulates cell population drug response over time

Next, to evaluate DREEP’s ability to recognize adaptation to drug treatment over time, we applied it to the Ben-David et al. dataset (66). As Figure 3A shows, this dataset comprises 7,440 cells from a single cell-derived clone of the MCF7 cell line exposed for two days to 500 nM of bortezomib, a 26S proteasome inhibitor. Specifically, cells were sequenced before treatment (t0), at 12 hrs and 48 hrs following drug exposure (t12, t48), and 96 hrs following drug washout and recovery (t96).

**Figure 3.**
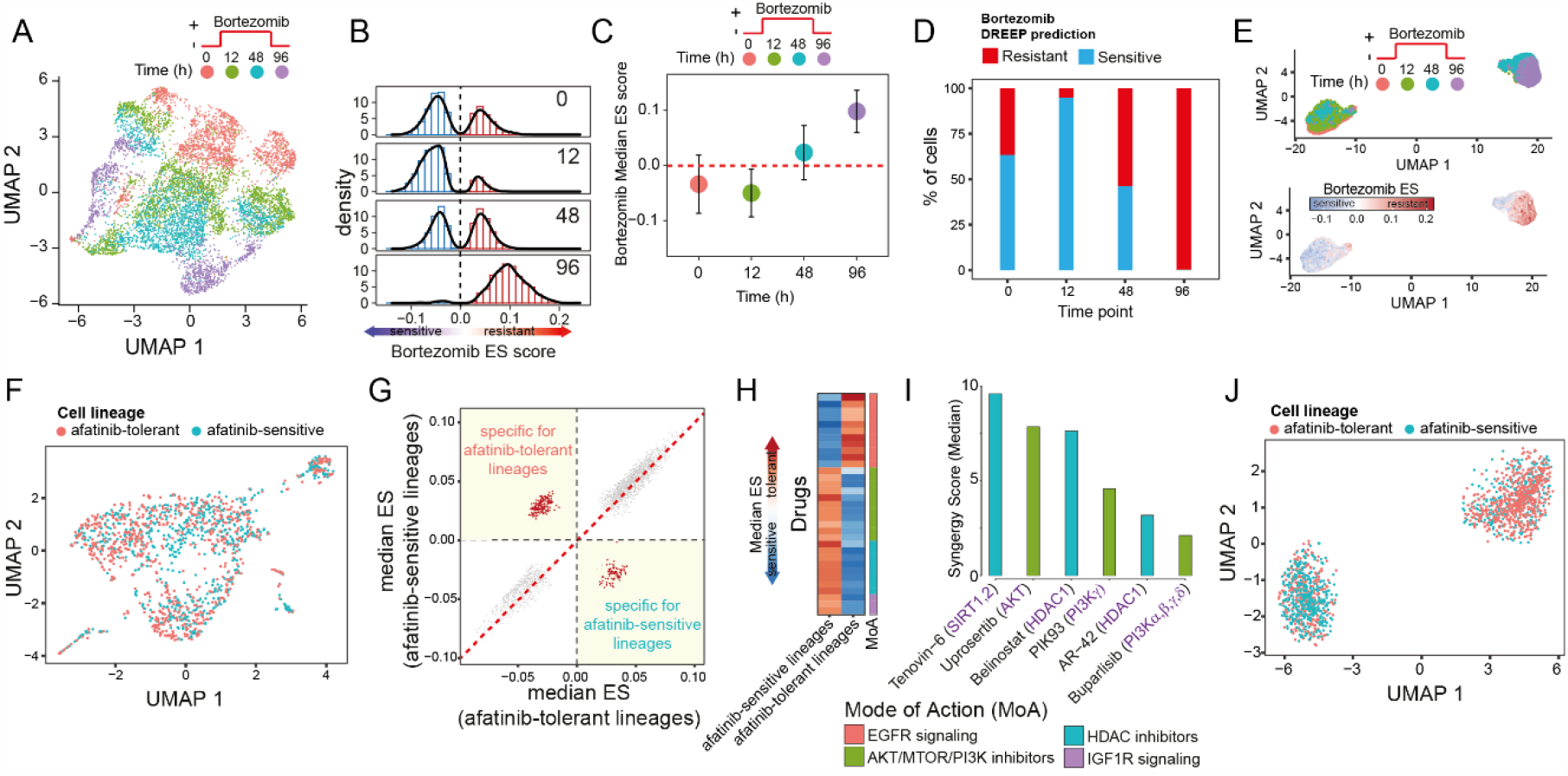
DREEP identifies drugs that can selectively inhibit a subpopulation of cells. (**A**) UMAP plot of 7,440 cells of the MCF7 cell line exposed to 500 nM bortezomib. (**B**) Bortezomib DREEP enrichment score (ES) distribution at each timepoint for the 7,440 cells of the MCF7 cell line in (A). A positive or negative score means that the cell is predicted to be resistant or sensitive to bortezomib, respectively. (**C**) Bortezomib DREEP’s median enrichment score of MCF7 cells at each time-point. (**D**) Percentage of predicted bortezomib sensitive or tolerant cells for each time point. (**E**) UMAP plot of the same cells in (A) but using the drug profile estimated by DREEP instead of its corresponding transcriptional profile for each cell. In the left panel, cells are colour-coded according to the sequencing timepoint, while in the right panel cells are colour-coded according to the bortezomib score predicted by DREEP. A positive or negative score means that the cell is predicted to be resistant or sensitive to bortezomib, respectively. (**F**) UMAP plot of 1,541 MDA-MB-468 cells from Pellecchia et al. (4). Cells are colour-coded according to whether they belong to an afatinib-sensitive or tolerant lineage. (**G**) Each point represents a drug whose coordinates are the median enrichment score predicted by DREEP for the drug in afatinib-tolerant cells [x-axis] and afatinib-sensitive cells [y-axis]. The more negative the value of the enrichment score, the more potent the predicted effect of the drug. The drugs specifically inhibiting at least one of the two subpopulations are indicated in red. (**H**) Heatmap showing enriched drug classes predicted to specifically inhibit afatinib-tolerant or afatinib-sensitive cells. Each row of the heatmap represents a drug coloured according to its median enrichment score predicted by DREEP across cells of the same lineage (i.e. afatinib-tolerant or afatinib-sensitive). (**I**) Median synergy score of drugs predicted by DREEP to selectively inhibit afatinib-tolerant lineages in MDA-MB-468 cells. Each drug was tested in combination with afatinib, and the synergy score computed using the Loewe statistical model. A positive score represents a synergism between the two tested drugs. (**J**) UMAP plot of the cells in (A) but using the drug profile estimated by DREEP instead of the corresponding transcriptional profile for each cell. Cells are colour-coded according to whether they belong to an afatinib-sensitive or tolerant lineage.

As shown in Figure 3B, the distribution of Bortezomib ES scores estimated by DREEP at time t0, t12 and t48 exhibits a bimodal pattern, indicating the presence of a heterogeneous cell population comprising both bortezomib-tolerant and bortezomib-sensitive cells (Figure 2E,F). By contrast, DREEP identified that at the final time point (t96), the distribution of bortezomib ES scores is unimodal, with a homogeneous cell population where most cells underwent transcriptome rewiring characterized by the expression of biomarker genes associated with bortezomib resistance (Figure 3C,D). Finally, we were interested in evaluating DREEP’s ability to detect distinct drug-response cell subpopulations before and after bortezomib treatment. To this end, we used drug response profiles estimated by DREEP instead of transcriptional profiles to build a new cell-embedding space (Methods). Figure 3E shows that the cell-embedding space built from DREEP sensitivity predictions is better than the mixed cell groups identified using transcriptional profiles (Figure 3A) at separating MCF7 cells into two main clusters resembling the drug exposure over time.

In summary, these analyses demonstrate DREEP’s ability to recapitulate drug response and variability over time.

### 5. The DREEP method identifies small molecules that selectively inhibit cancer cell subpopulations

Next, we evaluated DREEP’s ability to identify innate drug-resistant cell populations and to suggest drug co-treatments. To this end, we used single cell data of the untreated barcoded MDA-MB-468 breast cancer cell line that we recently generated (4). This dataset includes 1,541 single cells from the untreated MDA-MB-468 cell line, each associated with a molecularly expressed barcode (i.e., lineage). In the original study, the purpose of these barcodes was to enable single-cell lineage tracing within the MDA-MB-468 cell population. Subsequently, these cells underwent exposure to increasing concentrations of Afatinib, a tyrosine kinase inhibitor that targets EGFR and HER2 receptors, over an extended period exceeding 40 days. A bulk NGS approach was then used to identify the lineages (i.e., barcodes) that survived this prolonged Afatinib exposure. Finally, the surviving lineages were retrospectively mapped onto the single-cell sequencing data of the untreated barcoded MDA-MB-468 cells. This approach, known as “retrospective lineage tracing”, enabled the categorization of untreated MDA-MB-468 cells as either sensitive or tolerant to Afatinib and anti-EGFR treatments (4) (Figure 3F).

We applied DREEP to each of the two untreated MDA-MB-468 cell subpopulations (i.e. afatinib-tolerant and afatinib-sensitive) and predicted drugs able to selectively inhibit the growth of either. We thus found 156 drugs predicted to preferentially inhibit the afatinib-tolerant subpopulation and 80 drugs (FDR < 10% and Supplementary Table 05) for the afatinib-sensitive subpopulation (Figure 3G), which included 11 out of 12 EGFR inhibitors (Supplementary Table 05).

Interestingly, the most overrepresented classes among the drugs predicted to selectively inhibit the afatinib-tolerant subpopulation were IGF1R, HDAC, and AKT/MTOR/PI3K inhibitors (Figure 3H). We have already shown that afatinib and IGF1R inhibitors have a strong synergistic effect on these cells (4), thus validating the DREEP predictions that IGF1R inhibitors should deplete the cell population that do not respond to afatinib in this cell line.

To experimentally validate these predictions further, we selected three HDAC inhibitors (Tenovin-6, bellinostat, and AR-42) and three AKT/MTOR/PI3K inhibitors (Uprosertib, PIK93, and buparlisib) from the list of drugs predicted to selectively inhibit the afatinib-tolerant subpopulation. We then measured experimentally if these drugs had an additive, synergistic, or antagonistic effect on MDA-MB-468 cells in combination with afatinib (Supplementary Table 06). As shown in Figure 3I, the median synergy score of each of the six selected drugs tested in combination with afatinib is compatible with an additive effect (67), suggesting that the two drugs work independently on different subpopulations of cells. Finally, as Figure 3J shows, when cells are embedded into an embedding space reconstructed from DREEP sensitivity profiles (Methods), two main cell clusters appear that are enriched for either afatinib-tolerant or afatinib-sensitive lineages, in contrast to the mixed cell groups identified using transcriptional profiles (Figure 3F). These results show how the DREEP method alone can recognise that there are two different cell populations in this cell line characterized by a distinct drug response profile.

Altogether, these results show that DREEP can predict drug sensitivity at the single cell level and identify drugs able to selectively inhibit a subpopulation of cells that can be then combined to find drug combinations.

### 6. The DREEP method accurately predicts drug response in heterogeneous cancer patient cells

Cancer cell lines often display restricted transcriptomic heterogeneity, failing to mirror the full complexity of tumours. Therefore, to demonstrate the clinical applicability of DREEP, we utilized our methodology on a series of patient-derived cultures (PDCs) from both primary and metastatic sites of individuals suffering from head and neck squamous cell carcinoma (23). These models have already been proven to effectively replicate the complexity of the corresponding matching tumour (68), with the advantage that sensitivity measurements could be consistently assessed across multiple drugs (68). Specifically, this dataset comprises 1,027 single cell transcriptional profiles of PDCs obtained from five head and neck cancer patients whose drug response was measured in triplicate for five different small molecules at two different concentrations (18).

As illustrated in Figure 4A, the UMAP representation of single-cell transcriptomic profiles confirmed the presence of intra-patient transcriptomic heterogeneity within PDCs, particularly when juxtaposed with inter-patient heterogeneity. Notably, in line with observations previously reported in (23, 68), patients HN120 and HN137 displayed a significant degree of heterogeneity, where primary and lymph node metastatic tumour cells delineate two transcriptionally distinct cell subpopulations.

**Figure 4.**
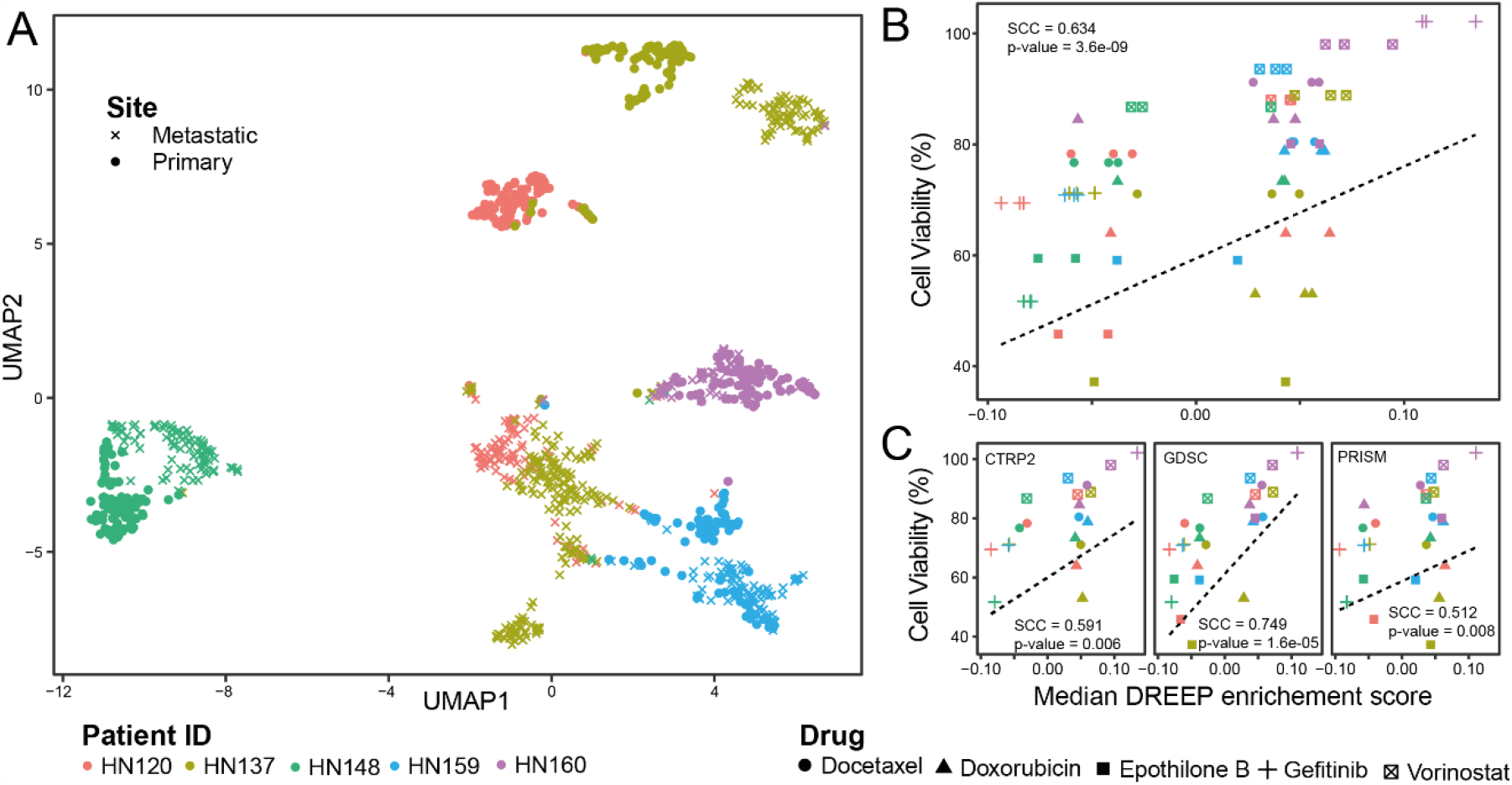
4 Drug response prediction in heterogenous patient-derived cultures (PDCs). (**A**) UMAP plot depicting 1,027 cells of PDCs from both primary and metastatic sites in five individuals with head and neck squamous cell carcinoma. (**B**) The Spearman correlation coefficient (SCC) between observed and predicted sensitivity scores for the five PDCs using five different drugs at lower concentrations. All three GDPS dataset are used. (**C**) A breakdown of (B) by GDPS datasets.

Next, to assess the predictive performance of DREEP on this dataset, we first applied it to the single-cell profiles of each patient and then converted these predictions from the single-cell level to the patient level by computing the median enrichment score for each drug across the cells within each individual patient. As depicted in Figure 4B, our analysis uncovered a significant correlation (Spearman’s correlation coefficient = 0.634, p-value = 3.6e-09) between DREEP predictions and the experimentally observed drug responses for five specific drugs: Docetaxel, Doxorubicin, Epothilone B, Gefitinib, and Vorinostat (refer to methods) (18). Patients’ cell viability was determined using each drug’s median IC50 values (18), which were threefold lower than those estimated from ATCC head and neck cancer cell lines present in the GDSC portal (methods). Finally, Figure 4C illustrates Spearman’s correlation coefficient (SCC) between DREEP predictions and the experimentally observed drug responses within each independent GPDS dataset. Notably, GDSC exhibited the strongest performance with an SCC of 0.749 (p-value = 1.6e-05), followed by CTRP2 with an SCC of 0.591 (p-value = 0.006), and PRISM with an SCC of 0.512 (p-value = 0.008). These findings remained consistent even when we employed patient cell viability estimates derived from drug concentrations matching the estimated IC50 values, as shown in Supplementary Figure 07.

## Discussion

Intra-tumour heterogeneity (ITH) presents a significant challenge in developing effective cancer treatments and achieving precision oncology. ITH refers to the variation of tumour cells both within and between tumours. As such, it is crucial to have methods to accurately identify the drug sensitivities of different tumour clones and determine the most effective treatment for each cancer type and the individual patient. To address this challenge, we introduced DREEP, a novel computational method that predicts drug sensitivity at the single cell level and can propose cell population-specific treatments. To achieve this, DREEP uses a set of drug-sensitivity signatures obtained by correlating gene expression and drug sensitivity profiles across hundreds of tumorigenic cell lines. It then uses these signatures to compute an enrichment score reflecting the degree of sensitivity of a cell to a specific drug.

To test the predictive performance and reliability of DREEP, we applied it to two independent datasets: (i) the pan-cancer cell line dataset from Kinker et al. comprising 198 tumorigenic cell lines from 22 distinct cancer types (30) and (ii) the single cell breast cancer atlas from Gambardella et al. that includes 32 breast cancer cell lines (5). We observed that DREEP successfully identifies the most effective drugs for each cell line with an overall performance that is several fold better than random on both datasets. We also successfully used DREEP to reconstruct the single-cell variability in drug response across the 198 cell lines of the pan-cancer dataset and to identify the functional cellular activities linked to common drug response patterns among cancer types and cell lines. Our approach can offer valuable insights for understanding the complex interplay between genetic backgrounds and drug sensitivity, paving the way for more personalized and effective cancer treatments. In addition, when used to analyse single-cell data collected from the breast cancer cell line MCF7 exposed to bortezomib over time (66), DREEP easily reconstructed the interchange between the sensitive and tolerant cell populations. Finally, using single cell data of the molecularly labelled MDA-MB-468 cell line (4), we showed that DREEP can identify innate drug-resistant cell populations and predict drug co-treatments able to deplete these cell populations. Overall, we show that DREEP produces an accurate reconstruction of the variability of drug response in cancer cell lines from single cell transcriptomics. The DREEP R package is available at https://github.com/gambalab/DREEP.

Our tool relies heavily on the quality and predictive power of GPDS, which were trained by linking gene expression to drug response data from pharmacogenomic screens. However, these screens may possess inherent biases and limitations in covering the broad spectrum of cancer types. We applied several evaluation metrics on multiple showcases to assess DREEP’s predictive accuracy and ability to generalize across different cancer types, encompassing approximately one hundred thousand cells originating from more than 20 distinct cancer types.

While our results have demonstrated DREEP’s ability to generalize across various cancer types effectively, we recognize that potential biases in our training data could limit DREEP’s applicability to underrepresented cancer types. Therefore, it is crucial to interpret DREEP’s performance within the context of the available training data. We advocate for ongoing refinement and expansion of pharmacogenomic screens to improve the diversity and coverage of cancer types. Addressing these biases in future data collection efforts can enhance DREEP’s utility in advancing personalized cancer treatment strategies. Moreover, in the future, it would be crucial to evaluate how the integration of other single-cell -omics measurements, such as genomic, proteomic, and epigenetic cell states, can improve the identification of variations in drug responses among cancer cell populations. Finally, since tumour heterogeneity is linked to clonal evolution processes, it would be also important to consider the dynamics of clonal populations under therapy pressure to improve the performance of future computational methods that will be developed for personalized diagnosis and treatment of cancer patients.

## Methods

### scRNA-seq datasets processing

The scRNA-seq data of the cell line pan-cancer dataset from Kinker et al. (30) was obtained from the Broad Institute’s single cell portal (SCP542). Specifically, CPM count matrix was downloaded. Before being processed, gene symbols were converted into ensemble id and only ensemble id associated with no more than one gene were retained. At the end of this pre-processing step, the final cell line pan-cancer matrix contained 16,558 genes and 53,513 cells. Raw scRNA-seq data of the MCF7 cell line exposed to bortezomib (66) was obtained from GEO (GSE114461) and only cells with at least 5,000 UMI were retained. The single cell breast cancer atlas dataset was obtained from figshare at the following address https://doi.org/10.6084/m9.figshare.15022698. scRNA-seq data of barcoded MDA-MB-468 cells were obtained from GEO (GSE228154). Raw UMI count data of nutlin treated and untreated cells was downloaded from figshare at https://figshare.com/s/139f64b495dea9d88c70 and only cells with at least 5,000 UMI and with less than 10% of UMI counts in mitochondrial genes were used. PDCs TPM counts were instead retrieved from supplementary data of the original work (18).

### Bulk RNA-seq dataset processing

Raw counts and associated metadata of the bulk RNA-seq data of cancer cell lines used to build GPDS signature were obtained from the depmap portal (https://depmap.org). Cell lines associated with haematopoietic tumours were excluded. Raw counts were normalised with edgeR (69) and transformed into log10 (CPM+1). Poorly expressed genes and genes whose entropy values were in the fifth percentile were discarded. Entropy values were estimated by discretizing the expression of each gene in ten equally spaced bins using the functions included in the R package named *entropy*. After these filtering steps, the final expression matrix comprised 1,117 cell lines and 13,849 genes.

### Drug sensitivity dataset processing

The drug sensitivity dataset of CTRP2 (Cancer Therapeutic Response Portal), GDSC (Genomics of Drug Sensitivity in Cancer) and PRISM (Profiling Relative Inhibition in Mixtures) were all obtained from the depmap portal (https://depmap.org). Only drugs for which the in vitro response was available for at least 100 cell lines out of the 1,117 for which gene expression data was available were retained. After this filtering, the final drug sensitivity dataset comprised 478 drugs from the GDSC dataset, 511 drugs from the CTRP2 dataset, and 1,445 drugs from the PRISM dataset. For all three cell viability datasets, drug potency was expressed in terms of AUC (Area Under the Curve) of the corresponding dose response graph.

### Construction of GPDS ranked-lists

Genomic Profiles of Drug Sensitivity (GPDS) ranked-lists were built by computing the Pearson correlation coefficient (PCC) between the expression of each gene and the potency of a drug across the cell lines for which drug potency was measured. At the end of this process, each GPDS is a ranked list of 13,849 expression-based biomarker genes whose degree of PCC reflects their importance in predicting the effect of a small molecule. This is because the AUC value of a drug reflects its cell line growth inhibition, with lower values indicating sensitivity and higher values implying tolerance to the drug. Thus, a gene whose expression positively correlates with the AUC of a drug is predictive of resistance to that drug, conversely a negatively correlated gene is predictive of sensitivity to the drug.

### Drug Response Estimation from single-cell Expression Profiles (DREEP)

To predict drug response at the single-cell level, we first used gf-icf normalization (https://github.com/gambalab/gficf) (27) to extract the top relevant genes from each cell. We then used these gene sets as input for Gene Set Enrichment Analysis (GSEA) (29) against each GDPS ranked-list to predict the sensitivity of a cell to a specific drug. Since GPDS ranked-lists contain predictive biomarkers of resistance at the top and predictive biomarkers of sensitivity at the bottom for each drug, a positive enrichment score implies that the cell expresses genes that render it tolerant to the drug, while a negative enrichment score indicates the cell expresses genes that render it sensitive to the drug. Finally, by estimating the median enrichment score on a cell population and counting the number of cells with a significant enrichment score less than zero, it is possible to estimate the overall effect of a drug on that population and the percentage of cells sensitive to it. The DREEP package is implemented in R and is available on github at the following address https://github.com/gambalab/DREEP. Enrichment Scores (ES) and associated p-values were computed using the *fgsea* package (70). The *fgsea* package utilizes an adaptive and efficient multi-level split Monte Carlo approach (71) to estimate the p-value of an ES. Subsequently, the p-values for each drug across the cells are corrected for the false discovery rate (FDR) using the Benjamini-Hochberg correction method, implemented within the function *p*.*adjust* in the *base* package of the R statistical environment. Finally, only ES values with an FDR < 0.1 were deemed significant (unless otherwise stated) and used to predict the effect of a drug on a single cell. The percentage of sensitive and resistant cells in a sample are finally computed using only significant ES scores.

### Estimation of DREEP performance

To estimate DREEP’s precision and sensitivity in predicting drug response from scRNA-seq data of cancer cell lines, we used the three cell-viability screening datasets from CTRP2, GDSC, and PRISM described above. For each cell-viability screening dataset, the corresponding “gold standard” is composed of pairs of cell lines and drugs for which the corresponding cell line is sensitive. To build the “gold standard” of each cell-viability screening dataset, we first computed the z-score percentiles from the AUC of each drug across the cell lines for which AUC was available and then defined a cell line sensitive to a drug only if its z-score was in the 5% percentile. Next, we applied DREEP to scRNA-seq data of each cell line and computed its median enrichment score for each drug. Finally, Precision-Recall and ROC curves on each cell-viability screening dataset depicted in Figure 1C-F were computed with the *precrec* R package (https://github.com/evalclass/precrec) using the corresponding “gold standard” and sorting DREEP drug/cell line pairs according to the estimated median cell line enrichment score multiplied by minus one in order to have drugs predicted to be sensitive at the top.

### Cell embedding space from DREEP predictions

DREEP output can be arranged in a drug response matrix of dimensions where is equal to the number of cells and equal to the number of GPDS used. Each element of matrix ranges within the interval and it is different from zero only if the enrichment score between the relevant genes of the cell and the GPDS ranked-list is significant (i.e. FDR < 0.1). Specifically, indicates the cell is predicted resistant to the drug, while indicates the cell is predicted sensitive to the drug and indicates that there is no supporting evidence to decide if cell is sensitive or not to the drug . Next, we computed the similarity of the predicted drug response profile between two cells as the distance between two cells and with *Fuzzy Jaccard Distance* defined as follow:

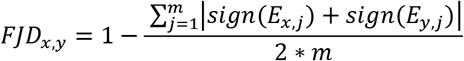

*FJD* is a non-negative symmetric distance matrix of dimensions whose elements range within the interval. Specifically, each element of quantifies how many of the drugs to which the cell is sensitive are shared with the cell and how many of the drugs the cell is resistant to being shared with the cell . The more two cells share drugs that are predicted to have the same effect, the closer the value is to 0, *and vice versa* for two cells with opposite drug predictions, equals 1. It can be easily shown that the Fuzzy Jaccard Distance we implemented satisfies all the conditions to be defined as a metric, including the triangle inequality constraint. Finally, the cell embedding space was built using the precomputed cell-to-cell distance matrix as input of the *umap* function of the uwot R package (https://github.com/jlmelville/uwot). Cell embedding from DREEP predictions has been implemented in the function *runDrugReduction* of the DREEP package.

### The therapeutic landscape of the cell line pan-cancer dataset (Figure 2A)

We first used DREEP to predict the effect of the 2,434 drugs on each of the 53,513 single-cell transcriptional profiles of the Kinker et al. dataset (30). Then, we converted predictions from the single-cell level to the cell-line level by counting the fraction of cells predicted to be sensitive to each drug (i.e. DREEP enrichment score < 0 and P < 0.05). At the end of this process, we obtained a matrix where contains the percentage of cells of the cell line predicted to be sensitive to the drug . Next, we set elements of less than 0.9 as being equal to 0 while all elements of greater or equal to 0.9 were set as equal to 1. Then, we hierarchically clustered rows (i.e. cell lines) and columns (i.e. drugs) of using the Jaccard distance. Jaccard distances between row pairs and column pairs of the matrix were computed using the *dist* function of the R statistical environment. Clustering of both, cell lines and drugs, was performed using the *hclust* function and resulting dendrograms cut with the *cutree* function of R statistical environment.

### Differential drug sensitivity analysis

We used DREEP to predict the effects of 2,434 drugs on 1,541 sequenced cells of the barcoded MDA-MB-468 cell line (4). Then, we performed a Mann-Whitney test and corrected p-values using the Benjamini-Hochberg method to identify drugs with differential sensitivity between afatinib-sensitive and afatinib-tolerant subpopulations. Finally, a drug was considered specific for a subpopulation if its FDR was less than 0.1 and the median enrichment score across the other subpopulation was greater than zero. This analysis has been implemented in the *runDiffDrugAnalysis* function of the DREEP package.

### Drug combinations assay

4 x 10^4^ MDA-MB-468 cells were seeded in a 96-well plate and incubated overnight at 37 °C. The two drugs to test were prepared in different dilutions and then combined in all possible drug pairs to generate a 5 x 5 combination matrix. Then, cells were exposed to either single agent drug or to the drug pairs of the drug combination matrix, while negative controls were treated with DMSO (each treatment was performed in triplicate). Following 72 hr incubation at 37 °C, cell viability was measured with CellTiter (Promega) and the absorbance was read at 490 nm with the plate reader GloMax® Discover instrument. The drug interactions and expected drug responses were calculated with the Combenefit tool (67) using the Loewe additivity model.

### Drug Set Enrichment Analysis (DSEA)

DSEA was employed by using the online drugEnrichr tool (31) using the Drug Repurposing Hub Mechanism of Action table resource.

### Comparison with other state-of-the-art methods

All methods were run using default parameters and when required single cell expression was normalized using the Seurat package. Common drugs between DREEP and each method were found by simply intersecting the drug names

### PDCs drug response data

PDCs drug response data were retrieved from supplementary data of the original work (18). Specifically, in the original work, eight drugs were tested on these five PDCs. However, we only used the five out of eight drugs for which cell viability was measured at least in triplicate. Raw viability measures we used can be found in Supplementary Table 07.

### Code availability

The DREEP package is available on github at the following address https://github.com/gambalab/DREEP.

### Data availability

All the processed datasets and associated metadata, including scRNA-seq, bulk RNA-seq, and sensitivity profiles used in this work have been deposited on figshare at the following address https://doi.org/10.6084/m9.figshare.23261234.

## Funding

This work was supported by the My First AIRC grant 23162.

### Author Contribution

SP and GV performed drug combination experiments. MF helped with bioinformatics analysis and in writing the manuscript. GG supervised the work, wrote the manuscript, and conceived the original idea.

### Competing interests

The authors declare no competing interests.

## Acknowledgement

We express our gratitude to Dr. Cathal Wilson for meticulously proofreading our manuscript. His invaluable efforts have significantly contributed to enhancing the clarity and accuracy of our work.

## Notes

https://github.com/gambalab/DREEP

### Competing Interest Statement

The authors have declared no competing interest.

### Summary of Updates

We added several new showcases after peer revision

