## Supplementary Figure 01 for "Predicting drug response from single-cell expression profiles of tumours" (from another version of this preprint: version 1, 2023-06-05)

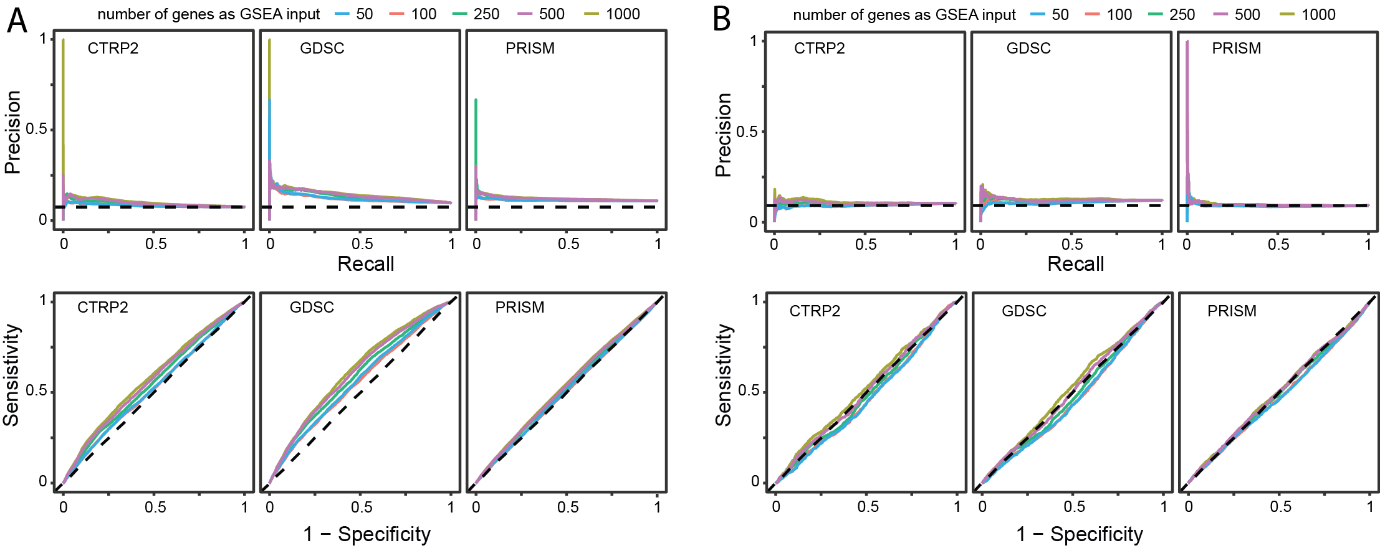

**Supplementary Figure 01 – Drug prediction performance using absolute expression to extract relevant genes of a cell.** (**A**) Performance of DREEP drug sensitivity on 198 cell lines for each set of GPDS signatures used and different numbers of most expressed genes to predict the effect of a drug. DREEP’s performance is estimated using either Precision-recall curve (upper) or ROC curve (bottom). (**B**) Same as (A) but for the 32 breast cancer cell lines published in Gambardella et al.
